# VSA-2, a novel plant-derived adjuvant for SARS-CoV-2 subunit vaccine

**DOI:** 10.1101/2025.09.22.677856

**Authors:** Awadalkareem Adam, Christy Lee, Madison C Jones, Brinley R Harrington, Jing Zou, Bi-Hung Peng, Xuping Xie, Pengfei Wang, Tian Wang

## Abstract

QS-21, a key component of several licensed vaccines is facing limited supply, dose-limiting toxicity and other drawbacks which together limit its broader usage. Development of saponin alternatives to QS-21 that retain its desirable adjuvant activity without its drawbacks is in high need. Incorporating an amide side chain into the more sustainable *Momordica* saponins (MS) I and II led to the recent discovery of two semisynthetic immunostimulatory adjuvants VSA-1 and VSA-2. Here, we showed that the receptor-binding protein (RBD) of ancestral SARS-CoV-2 adjuvanted with VSA-2 (VSA-2-RBD) induced high titers of SARS-CoV-2-specific humoral and T helper-1 prone immune responses in mice comparable to that triggered by QS-21-RBD. Vaccination with VSA-2-RBD provided strong protection against SARS-COV-2, Delta and Omicron variants infection and the virus-induced lung inflammation and pathology similarly as QS-21-RBD vaccination. Overall, our results suggest that VSA-2 adjuvant can potentially complement the clinically proven saponin adjuvant QS-21 in vaccines against infectious diseases.

## 1. Introduction

QS-21, a saponin isolated from the tree bark extract of *Quillaja saponaria* (QS) Molina, is known to induce strong and balanced humoral and cellular immune responses. It has been used as an adjuvant in licensed and exploratory vaccines and preclinical studies[1-6]. QS-21 in combination with the toll-like receptor (TLR) agonists, such as Monophosphoryl lipid A (MPL) and/or CpG are also the key components of several adjuvants, including AS01 (QS-21 and MPL), AS05 (QS21, MPL, and alum), and AS15 (QS-21, MPL, and CpG) [1, 5, 7-9]. Despite its success in enhancing safety, efficacy and durable vaccination, QS-21 has been reported to have several drawbacks, such as limited supply, dose-limiting toxicity, laborious and low-yielding purification and chemical instability which together limit its broader usage[10-12]. Development of saponin alternatives to QS-21 that retain its desirable adjuvant activity without its drawbacks is highly desirable, and is more advantageous over developing other classes of vaccine adjuvants[13].

*Momordica* saponins (MS) I and II are more sustainable and share high resemblance to the deacylated QS saponins like QS-21. Incorporating an amide side chain into the MSI and II, led to the recent discovery of two MS-derived semisynthetic immunostimulatory adjuvants, VSA-1 and VSA-2[14, 15]. The VSA adjuvants are much more accessible and of lower toxicity compared to QS-21. VSA-1 was reported to stimulate similar or higher levels of antibody titers and provided protection upon vaccination with pneumococcal vaccine PCV13 or inactivated influenza virus compared to those by QS-21[16, 17]. Although VSA-2 was found to induce comparable levels of antigen-specific IgG and subtype antibodies and T cell responses upon vaccination with ovalbumin antigen than that by QS-21-adjuvanted vaccination[18], limited data are currently available on VSA-2 -adjuvating activity in boosting subunit vaccines of infectious agents.

Severe acute respiratory syndrome coronavirus 2 (SARS-CoV-2) is the causative agent of the recent COVID-19 pandemic. The virus is a member of the genus *Betacoronavirus* (β-CoV) and the family *Coronaviridae*[19]. The single-stranded positive-sense RNA genome encodes structural proteins (spike [S], envelope [E], membrane [M], and nucleocapsid [N]), nonstructural proteins (nsp1-nsp16), and several accessory proteins [20]. Among them, the S and its receptor-binding domain (RBD) are often the main targets for vaccine development due to its roles in facilitating virus entry into target cells via interactions with the human angiotensin-converting enzyme 2 (hACE2) receptor and its enrichment of T and B cell epitopes[21-23]. Several major vaccine platforms which were approved by the FDA for Emergency Use Authorizations (EUAs) during the pandemic are now licensed for use, including the mRNA vaccines from Moderna (Spikevax) and Pfizer-BioNTech (Comirnaty) and the subunit protein vaccines Novavax (Nuvaxovid) [24]. Despite their roles in successful reduction of the virus-induced disease severity, hospitalizations, and mortality during the pandemic, further optimizing existing vaccine platforms and developing more effective novel vaccines are needed to reduce the safety concerns of exposure to continuously circulating variants post-pandemic. Furthermore, QS-21 is the main component of Matrix-M, the adjuvant system used in the Nuvaxovid [25, 26]. Thus, in this study, we evaluated VSA-2 adjuvanted SARS-CoV-2 RBD subunit vaccine (VSA-2-RBD) in boosting humoral and cell-mediated immunity in mice and assessed the protective efficacy of VSA-2-RBD vaccination against SARS-CoV-2 and variants infection.

## 2. Results

### Vaccination with VSA-2-RBD subunit vaccine induced potent T helper 1 (Th)1 and Th2-prone humoral and cell-mediated immune responses in mice

To assess the adjuvant activity of VSA-2 for the SARS-CoV-2 subunit vaccines, BALB/c mice were subcutaneously (s.c.) inoculated with PBS (mock), QS-21-RBD (10 µg), or VSA-2-RBD (10 µg) on day 0 and boosted with the same dose on day 21. Sera were collected at 31 days post vaccination (DPV) to determine antibody titers (**Fig. 1A**). VSA-2-RBD-vaccinated mice showed strong RBD-binding IgG antibody responses, which were comparable to that of the QS-21-RBD-vaccinated mice (**Fig.1B-C)**. The production of IgG1 and IgG2a antibodies is often used as an indication for Th1 and Th2 prone responses respectively as cytokines produced by these T cell subsets mediate respective isotype switching[27]. VSA-2-RBD vaccination induced about 6 to 7 logs titers of SARS-CoV-2 -specific IgG2a and IgG1 antibodies at 31 DPV (**Fig. 1D-E**). Similar titers of IgG2a and IgG1 antibodies were detected in the QS-21-RBD group. No significant differences were noted between the two vaccination groups. Thus, vaccination with VSA-2-RBD induces potent Th1 and Th2-prone SARS-CoV-2-specific antibody responses comparable to the QS-21-RBD vaccination.

**Figure 1.**
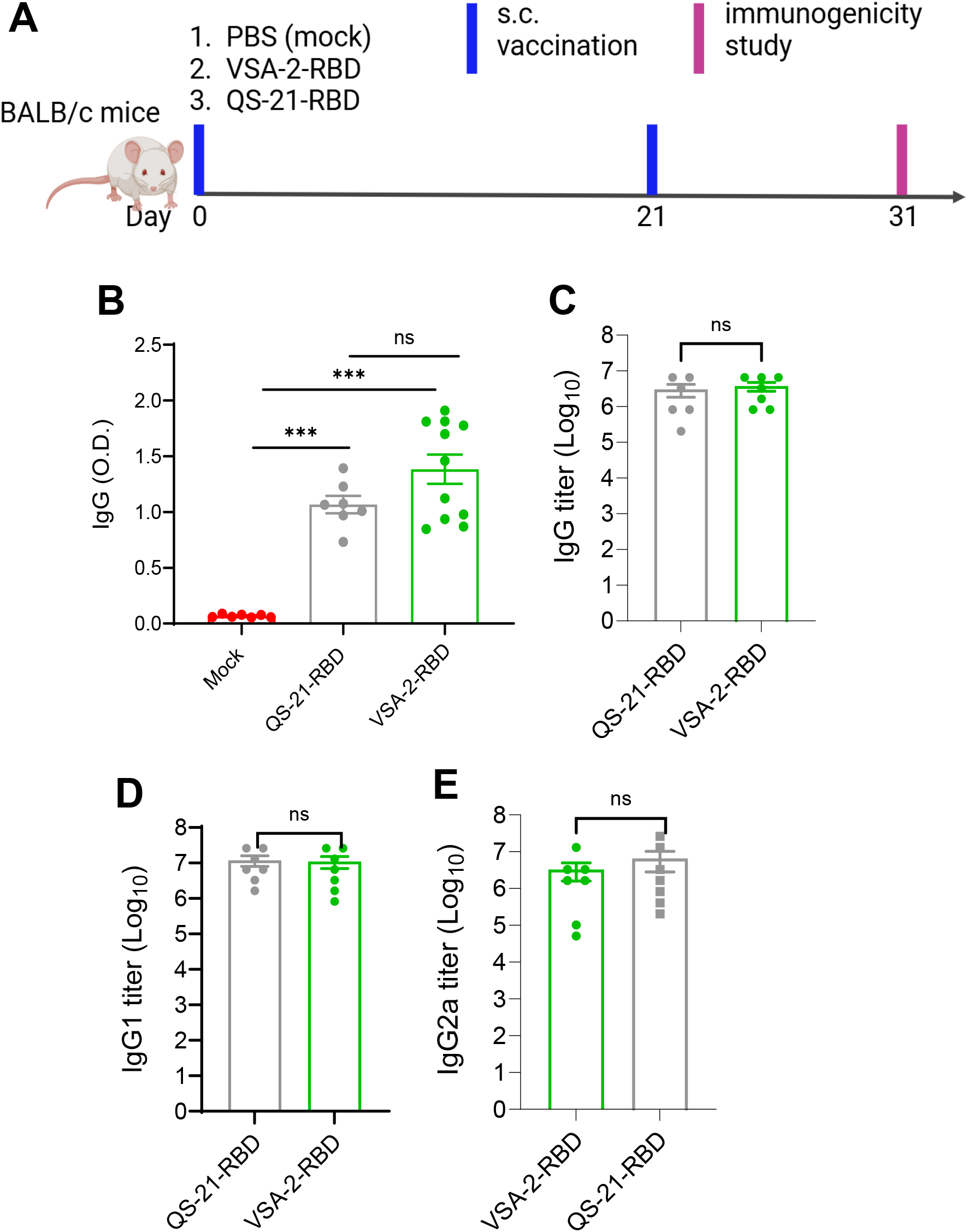
VSA-2 RBD vaccination induced strong Th1 and Th-2 prone SARS-CoV-2 IgG binding antibodies. BALB/c mice were prime-boost immunized with mock (PBS), QS-21-RBD, or VSA-2-RBD via s.c. route. **A**. Study design. **B-C**. ELISA O.D. values and the endpoint IgG titers against SARS-CoV-2 RBD measured in serum. n = 6 to 11. **D-E**. Endpoint IgG subtype titers against SARS-CoV-2 RBD measured in serum. *** *P* < 0.001 compared to the mock group.

Next, to assess the VSA-2-RBD-induced T cell immunity, spleen samples were collected on 31 DPV and the splenocytes were stimulated *in vitro* with SARS-CoV-2 S peptide pools. ELISPOT analysis showed that both VSA-2-RBD and QS-21-RBD groups had more than 40-fold higher IFN-γ production compared to the mock group (**Fig. 2A-B**). No differences were noted between the VSA-2-RBD and QS-21-RBD groups. We further analyzed CD4^+^ and CD8^+^ T cell responses in the vaccinated mice. At 31 DPV, the percent and total cell number of CD4^+^IFNγ^+^ T cells in the VSA-2-RBD-vaccinated mice were about 2.5-fold higher than that of the mock group; whereas CD4^+^IFNγ^+^ T cells of the QS-21-RBD-vaccinated mice increased 1.3-fold only by cell number (**Fig. 2C-D**). Furthermore, both the VSA-2-RBD and QS-21-RBD groups showed an induction of CD8^+^IFNγ^+^ T cells by percent and total cell number; the QS-21-RBD group had a modest higher increase in cell number than VSA-2-RBD group (**Fig. 2E-F**). Overall, these results suggest that VSA-2-RBD vaccination induces strong Th1 and Th2 prone antibody and cell-mediated immune responses comparable to that of QS-21-RBD vaccination. VSA-2 -RBD triggered higher CD4^+^IFNγ^+^ T cell responses.

**Figure 2.**
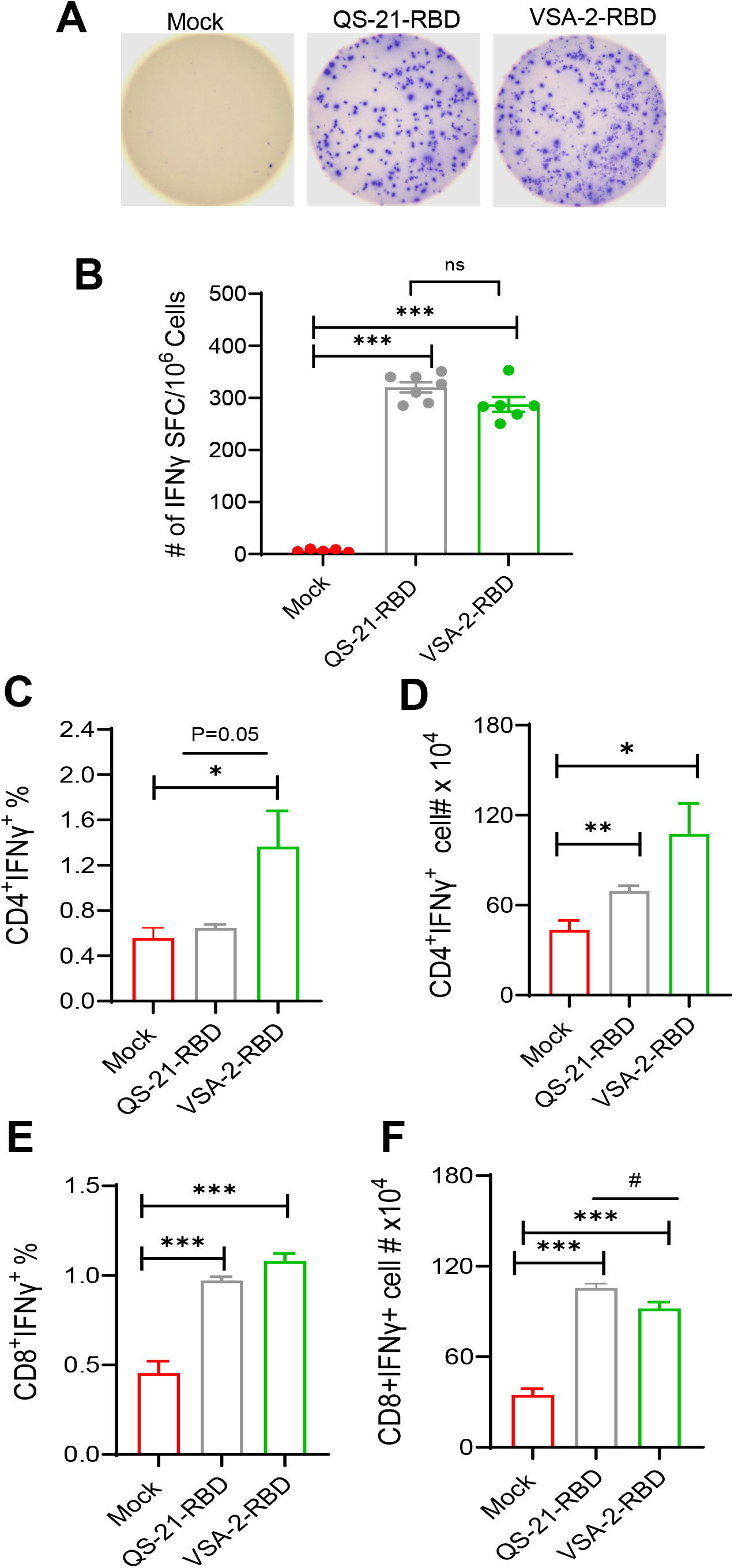
VSA-2 RBD vaccination induced potent Th-mediated immune responses. BALB/c mice were prime-boost immunized with mock (PBS), QS-21-RBD, or VSA-2-RBD via s.c. route. **A-B**. ELISPOT quantification of vaccine-specific splenic T cells at 31 DPV. Splenocytes were stimulated with SARS-CoV-2 S peptides, or blank for 20 h. **A**. Images of wells from T cell culture. **B**. Spot forming cells (SFC) were measured by IFN-γ ELISpot. Data are shown as # of SFC per 10^6^ splenocytes. n= 5-7. **C-G**. Splenocytes were stimulated with SARS-CoV-2 S peptides *ex vivo* for 6 h, and stained for IFN-γ, CD3, CD4, or CD8. Percent and total number of IFN-γ^+^ CD4^+^ and CD8^+^ T cells are shown. \*\*\**P* < 0.001, \*\**P* < 0.01, or \**P* < 0.05 compared to mock. ^#^*P* < 0.05 compared to QS-21-RBD group.

### Vaccination with VSA-2-RBD provides strong protection against SARS-COV-2 and the variants infection as well as the virus-induced lung inflammation and pathology

To evaluate the protective efficacy of VSA-2-RBD vaccination against SARS-CoV-2 infection, BALB/c mice were vaccinated with mock, VSA-2-RBD (10 µg), or QS-21-RBD (10 µg) s.c. on day 0 and boosted with the same dose on day 21. At 31 DPV, mice were i.n. challenged with 2 x 10^4^ PFU mouse-adapted SARS-CoV-2 strain CMA4 [28]. On day 2 post-infection, lung tissues were collected to measure viral loads and inflammatory immune responses and histopathology to evaluate the protective efficacy (**Fig. 3A**). Plaque and Q-PCR assays suggest significantly diminished levels of viral RNA in the lungs of both VSA-2-RBD and QS-21-RBD groups and barely any detectable viral particles (**Fig. 3B-C**). No differences were noted between the two groups. Sera from the vaccinated mice were also collected on 31 DPV to test for neutralization antibody titers against SARS-CoV-2 Delta and Omicron BA.2 variants (**Fig. 3D-E**). Both VSA-2-RBD and QS-21-RBD groups showed 3 to 4 logs of neutralization antibody titers against the SARS-CoV-2 Delta variant. The VSA-2-RBD group had about 2 to 3 logs of neutralization antibody titers against the B.A.2 variant with a trend slightly higher than the QS-21-RBD group.

**Figure 3.**
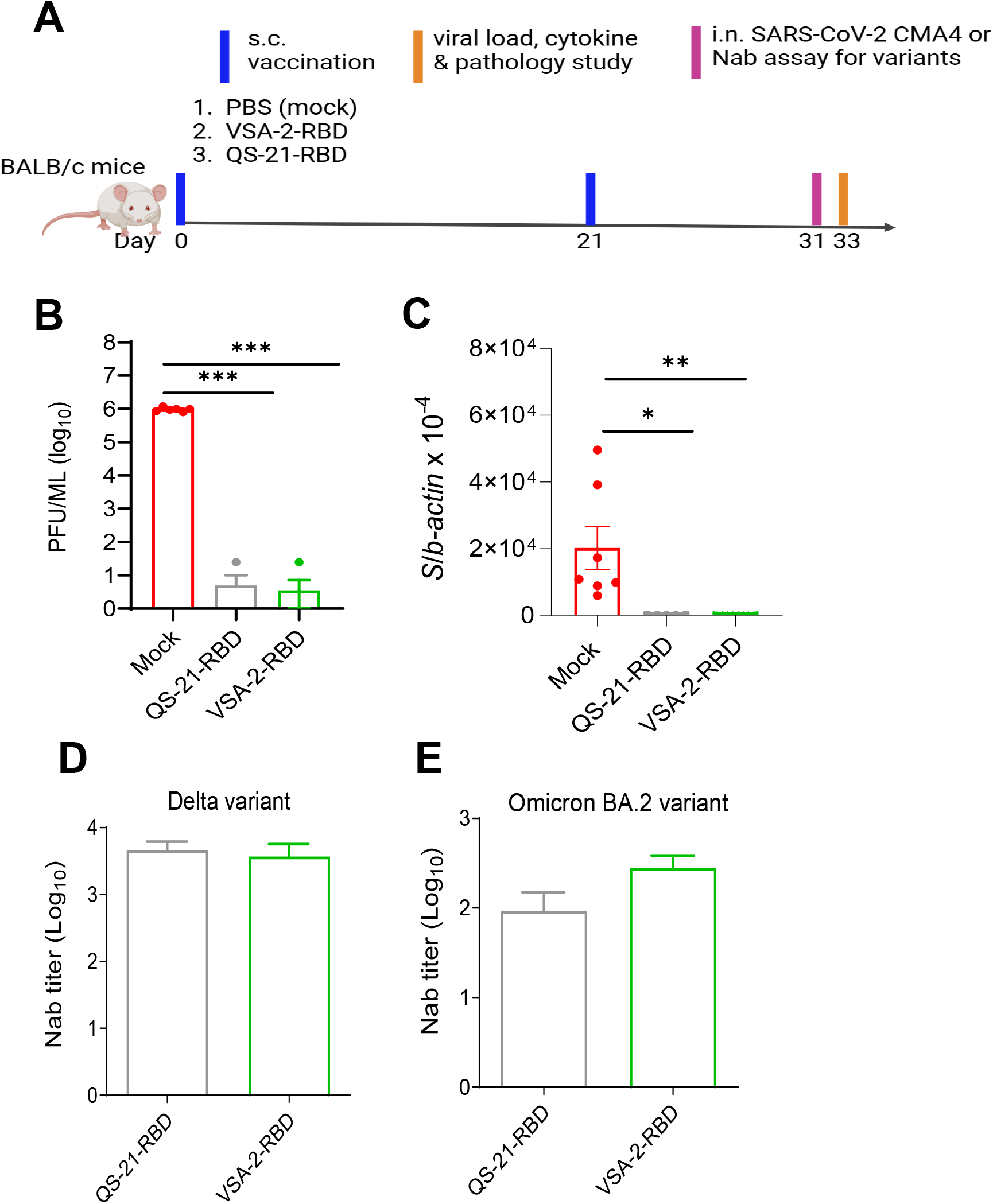
Vaccination with VSA-2-RBD protected mice from SARS-coV-2 and variants infection. BALB/c mice were prime-boost immunized with mock (PBS), QS-21-RBD, or VSA-2-RBD via s.c. route. At 31 DPV, mice were i.n. challenged with 2 x 10^4^ PFU mouse-adapted SARS-CoV-2 strain CMA4. On day 2 post infection, lung tissues were collected to measure viral loads. **A**. Study design. **B-C**. Viral loads were measured by plaque or Q-PCR assays. n= 5 to 7. **D-E**. Serum neutralizing activity against mNG SARS-CoV-2 Delta and Omicron BA.2 variants was measured by plaque reduction neutralization test (PRNT). mNG-NT_50_ titers are shown, n = 3 to 5. Data are presented as means ± SEM. \*\*\**P* < 0.001, \*\**P* < 0.01, or \**P* < 0.05 compared to mock.

As SARS-CoV-2-induced cytokine storm in the lung is known to be associated with COVID-19 disease severity in humans [29, 30], we next determined proinflammatory cytokine and chemokine production in murine lungs at day 2 post challenge by Q-PCR. We observed that both VSA-2-RBD and QS-21-RBD groups showed significantly lower levels of IL-6, CCL2, CCL7 and CXCL10, whereas a significant reduction on IL-1β expression was only observed in the VSA-2-RBD group (**Fig. 4A-E**). Furthermore, lung histopathology examination (**Fig. 4F**) reveals that the mock group had severe respiratory epithelial detachment/death without recruiting much of inflammatory cells to the affecting areas. In comparison, the VSA-2-RBD and QS-21-RBD groups showed few small foci of respiratory epithelial detachment with minimal peri-bronchial infiltrations but had heavier peri-vascular lymphocytic infiltrates, especially in VSA-2-RBD group as shown in **Fig. 4F**, which indicate VSA-2-RBD-induced T cell responses are involved in host protection upon SARS-CoV-2 challenge. In summary, these data suggest that VSA-2-RBD and QS-21-RBD prime-boost vaccination protects mice from viral infection and virus-induced inflammatory responses and pathology in the lung by induction neutralization antibodies and T cell-mediated immunity.

**Figure 4.**
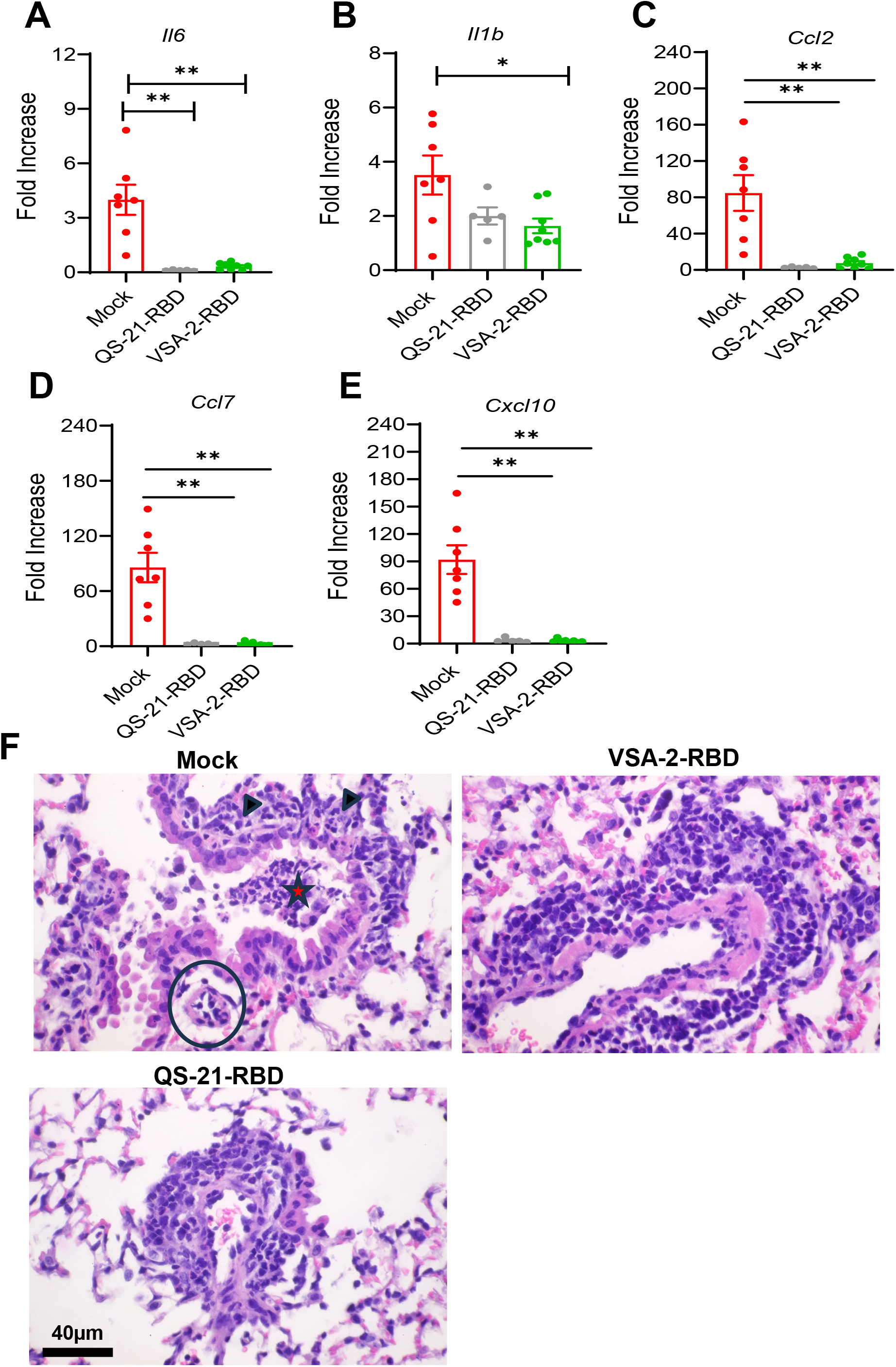
Vaccination with VSA-2-RBD protected mice from SARS-coV-2 -induced lung inflammation and pathology. BALB/c mice were prime-boost immunized with mock (PBS), QS-21-RBD, or VSA-2-RBD via s.c. route. At 31 DPV, mice were i.n. challenged with 2 x 10^4^ PFU mouse-adapted SARS-CoV-2 strain CMA4. At day 2 post infection, lung tissues were collected. **A-E**. Lung cytokine and chemokine levels at day 2 post infection determined by Q-PCR assays. Data is presented as the fold increase compared to mock-infected control. **F**. Lung sections were prepared for histopathology examination. Mock group (top left panel) showed widespread respiratory epithelial detachment (red star) with mild peri-bronchial infiltrations with mononuclear cells (arrowheads), and minimal in peri-vascular space (circle). QS-21-RBD (low left panel) & VSA-2-RBD group (top right panel) induced higher levels of lymphocytic infiltrates but less epithelial injury. Bar = 40 µm. Data are presented as means ± SEM. ** *P* < 0.01 or * *P* < 0.05 compared to mock group.

## 3. Discussion

Balanced humoral and Th-1 directed cellular immune responses are important for host protection against SARS-CoV-2 infection[31]. QS-21, a key adjuvant component of several licensed vaccines [1-6], has been shown to efficiently adjuvant SARS-CoV-2 subunit vaccines in boosting protective immunity [32-34]. In this study, we demonstrate that vaccination with VSA-2, a recently discovered MS-derived semisynthetic immunostimulant, together with the SARS-CoV-2 RBD antigen induced both Th1 and Th2 prone antibody responses and T cell-mediated immune responses to the levels comparable to those triggered by QS-21-RBD vaccination in mice. In addition, VSA-2-RBD vaccination protected mice from SARS-CoV-2 infection and the virus-induced lung inflammation and pathology. VSA-2 RBD vaccination also triggered strong neutralization antibodies against SARS-CoV-2 Delta and Omicron B.A.2 variants. In summary, our results suggest that VSA-2 displays an adjuvant activity similar to QS-21 in SARS-CoV-2 subunit vaccination.

Prior studies suggest a dose of 10 to 20 µg of QS-21 in combination with other adjuvants would achieve robust immune responses in vaccination[35-37]. Here, we noted that vaccination with 10 and 15 µg of QS-21-combined with the RBD antigen induced potent SARS-CoV-2 IgG antibody response with no significant differences between the two doses (**Supplementary Fig.1**). Thus, 10 µg of QS-21 was used in prime and boost vaccination of this study. A dose of 100 µg of VSA-2 was previously shown to be optimal for its adjuvant activity [38]. The same dose was also used in this study. Further studies will be optimized to determine the minimal dose of VSA-2 for protection and strong immunogenicity. Furthermore, liposome is known to act as delivery vehicles to encapsulate antigens and adjuvants and improve the uptake by antigen-presenting cells (APCs). Liposome formulation of QS-21 was reported to reduce the dosage for both QS-21 and the antigen with enhanced immune responses[39]. Indeed, liposomated QS-21 together with the SARS-CoV-2 spike ferritin nanoparticle vaccine triggered enhanced antigen-specific humoral and T cell responses [40]. Liposome formulation of VSA-2 may be tested in future optimization studies of the adjuvant.

QS-21 in combination with the TLR agonists, including MPL and/or CpG are key components of several licensed adjuvants, including AS01, AS05, and AS15 [5, 7-9] and were shown to more efficiently boost host humoral and cellular immune responses during vaccination. For example, QS-21 together with MPL acted synergistically in enhancing cell-mediated immune responses in vaccination of varicella-zoster virus glycoprotein [41]. The combination of QS-21 with MPL or CpG was also shown to enhance antigen-specific polyfunctional T cell response, virus-neutralization antibody and IgG binding antibody responses with broad immunogenicity during SARS-CoV-2 subunit protein vaccination in mice, and non-human primates [32, 42, 43]. Optimization of VSA-2 /TLR agonists combination adjuvant is underway and will continue to be the focus of our future investigation. The results will likely provide mechanistic insights for rational design of new adjuvants with optimal properties.

Intranasal immunization can lead to the induction of antigen-specific immunity in both the mucosal and systemic immune compartments [44]. Nevertheless, currently licensed SARS-CoV-2 vaccines, including the subunit vaccines are limited to the parenteral injection. One of the challenges is that soluble antigens delivered to the nasal passages do not breach the epithelial barrier but instead are transported by microfold cells [45]. We previously reported that a modified porous silicon microparticle (mPSM) facilitates mucosal delivery of antigen to the mucosal sites. Parenteral and intranasal prime and boost vaccinations with mPSM-RBD triggered robust mucosal and systemic immunity [46]. The liposome formulation of VSA-adjuvant makes it easy for mucosal delivery of antigens, which would allow testing the immunogenicity and protective efficacy of the hybrid vaccination platform in future vaccine development.

The use of recombinant protein subunits as the candidate vaccine antigens is considered to be a safe and reliable technique. Compared to the currently available adjuvants, VSA-1 induced higher or equivalent levels of antibody titers and enhanced protection following vaccination with the influenza virus and pneumococcal glycoprotein compared to those by Q-21 adjuvant[16, 17]. Our study and two previous reports demonstrate that VSA-2 induces broad, stronger or equivalent humoral and cellular immune responses [18, 47]. Thus, the Momordica saponin-derived VSA adjuvants will address the supply shortage of QS-21 and can complement QS-21 with different adjuvant properties to meet the requirements of various vaccine applications. The VSA adjuvants will also facilitate the development of various chemical combinations of adjuvants and built-in adjuvants, thereby enabling the creation of multicomponent synthetic vaccines. In summary, VSA adjuvants represent a new generation of saponin adjuvants that can potentially complement the clinically proven saponin adjuvant QS-21 for broader applications in vaccines against infectious diseases.

## 4. Materials and methods

### Ethic Statement

All animal experiments were approved by the Animal Care and Use Committees at the University of Texas Medical Branch (UTMB) (protocol #1412070).

### General procedure of derivatizing MS II

Momordica saponins were isolated by using the published procedure[48] To a clear solution of MS II (15.8 mg, 9.4 µmol) in ethanol (0.5 mL) and water (drops) was added 11-aminoundecanoic acid benzyl ester hydrochloride (6.6 mg, 20 µmol),[49] *N*-methylmorpholine (NMM) (12.0 mg, 118 µmol), hydroxybenzotriazole (HOBt) (9.2 mg, 60 µmol), and 1-ethyl-3-(3-dimethylaminopropyl)carbodiimide hydrochloride (EDC.HCl) (12.0 mg, 63 µmol) at room temperature.[50] The reaction mixture was stirred for 1 day and then filtered. The filtrate was purified with RP HPLC by using a semi-Prep C18, 250×10 mm, 5 micron column and H_2_O/MeCN gradients (90%-10% H_2_O over 45 minutes with a 3 mL/min flow rate). The desired product had a retention time of 30 min and the fraction was concentrated on a rotary evaporator at room temperature to remove MeCN, and the remaining water was then removed on a lyophilizer to provide final product VSA-2 as a white solid.

### Vaccination and challenge of mice

Six to seven-week-old BALB/c mice were purchased from The Jackson Laboratory. For vaccination, mice were injected subcutaneously (s.c.) with 10 µg RBD (Ray Biotech) mixed with 10 or 15 µg QS-21 (InvivoGen), 100 µg VSA-2 (??), or PBS (mock) on days 0 and 21. For challenge studies, vaccinated mice were infected intranasally (i.n.) with 1×10^4^ plaque-forming units (PFU) of a mouse-adapted SARS-CoV-2 CMA4 strain. Infected mice were monitored daily for signs of morbidity. On days 2 post-infection, mice were euthanized for tissue collection. The right superior lobes of lung tissues were collected in Trizol for RNA extraction for viral load and cytokine/chemokine analysis. The inferior lobes of the right lung were homogenized in 1 ml of DMEM for viral titration by plaque assay.

### Antibody ELISA

ELISA plates (Corning, USA) were coated with 100 ng/well recombinant SARS-CoV-2 RBD protein (Ray Biotech) for overnight at 4°C as described previously [46, 51]. The plates were washed twice with PBS containing 0.05% Tween-20 (PBS-T) and then blocked with 8% FBS for 1.5 h. Sera were diluted 1:100 or 1: 2 serial dilutions in blocking buffer. Samples were added for 1 h at 37°C. Plates were washed 5 times with PBS-T. For IgG measurement, goat anti-mouse IgG (Southern Biotech) coupled to peroxidase (HRP) was added at a 1:2000 dilutions for 1 h at 37ºC followed by adding TMB (3, 3, 5, 5′-tetramethylbenzidine) peroxidase substrate (Thermo Fisher Scientific) for about 5 min. The reactions were stopped by 1M sulfuric acid, and the intensity was read at an absorbance of 450 nm. For IgG1 and IgG2a titration, goat anti-mouse IgG1 or IgG2a (both from Southern Biotech) coupled to alkaline phosphatase (ALP) was added at 1: 2000 dilutions for 20 min and the plate were read at an absorbance of 405 nm. Binding endpoint titers were determined using a cutoff value which is negative control +3 x SD.

### IFN-γ ELISPOT

Millipore ELISPOT plates (Millipore Ltd) were coated with mouse anti-IFN-γ capture Ab at 1:100 dilution (Cellular Technology Ltd) and incubated at 4°C overnight. Splenocytes cells were stimulated with SARS-CoV-2 S peptide pool (1 μg/ml, Miltenyi Biotec) for 24 h at 37°C. Cells were stimulated with anti-CD3 (1 μg/ml, e-Biosciences) or medium alone were used as controls. This was followed by incubation with biotin-conjugated anti-IFN-γ at 1:100 dilution (Cellular Technology Ltd,) for 2 h at room temperature, followed by incubation with alkaline phosphatase-conjugated streptavidin for 30 min. The plates were washed and scanned using an ImmunoSpot 6.0 analyzer and analyzed by ImmunoSpot software to determine the spot-forming cells (SFC) per 10^6^ lung leukocytes.

### Intracellular cytokine staining (ICS)

Splenocytes were incubated with SARS-CoV-2 Spike protein peptide pools (1μg/ml, Miltenyi Biotec) for 6 h. BD GolgiPlug (BD Bioscience) was added to block protein transport at the final 1 h of incubation. Cells were stained with antibodies for CD3 (12-0031-81), CD4 (17-0041-82), or CD8 (11-0081-82) purchased from Thermo Fisher Scientific, fixed in 2% paraformaldehyde (PFA), and then permeabilized with 0.5% saponin before adding anti-IFN-γ (12-7311-82, Thermo Fisher Scientific). Samples were processed with a C6 Flow Cytometer instrument. Dead cells were excluded based on forward and side light scatter. Data were analyzed with a CFlow Plus Flow Cytometer (BD Biosciences).

### Fluorescent focus reduction neutralization test

Neutralization titers of mice sera were measured by a fluorescent focus reduction neutralization test (FFRNT) using the mNG reporter viruses SARS-CoV-2 Delta and BA.2 as previously reported with some modifications [52]. Briefly, Vero E6 cells (2.5 × 10^4^) were seeded in each well of black μCLEAR flat-bottom 96-well plate (Greiner Bio-one™). The cells were incubated overnight at 37°C with 5% CO_2_. On the next day, each serum was 2-fold serially diluted in the culture medium with the first dilution of 1:10. Each serum was tested in duplicates. The diluted serum was incubated with 100-150 fluorescent focus units (FFU) of mNG Delta or BA.2 SARS-CoV-2 at 37°C for 1 h (final dilution range of 1:20 to 1:20480), after which the serum-virus mixtures were inoculated onto the pre-seeded Vero E6 cell monolayer in 96-well plates. After 1 h infection, the inoculum was removed and 100 μl of overlay medium (DMEM supplemented with 0.8% methylcellulose, 2% FBS, and 1% P/S) was added to each well. After incubating the plates at 37 °C for 16 h, raw images of mNG fluorescent foci were acquired using CytationTM 7 (BioTek) armed with 2.5× FL Zeiss objective with wide field of view and processed using the software settings (GFP [469,525] threshold 4000, object selection size 50–1000 µm). The foci in each well were counted and normalized to the non-serum-treated controls (set as 100%) to calculate the relative infectivity. The neutralizing titer 50 (NT_50_) was calculated manually as the highest dilution of the serum sample that prevents at least 50% fluorescence foci formation in infected cells. A titer is calculated for each of the two replicates of a sample and the geometric mean titer (GMT) of the two is reported as the final sample titer.

### Quantitative PCR (Q-PCR)

Lung tissues were resuspended in TRIzol for RNA extraction according to the manufacturer’s instruction (Life Technologies). RNA concentration and purity was determined using a WPA Biowave DNA Spectrophotometer. Complementary (c) DNA was then synthesized with a qScript cDNA synthesis kit (Bio-Rad). Expression of SARS-CoV-2 Spike gene and mouse cytokine and chemokine genes (IL-1β, IL-6, CCL2, CCL7, and CXCL10) was measured by qPCR using the CFX96 real-time PCR system (Bio-Rad). PCR cycling conditions were as follows: 95°C for 3 min, 45 cycles of 95°C for 15 s, and 60°C for 1 min. Gene expression was calculated using the formula 2^ -[Ct(target gene)-Ct(β-actin)], as described previously [53].

### Plaque assay

Vero E6 cells were seeded on 6-well plates and incubated at 37°C for 16 h. Lung tissue homogenates were subjected to 10-fold serial dilutions in DMEM with 2% FBS and 0.2 ml dilutions were used to infect cells for 1 h. After the incubation, samples were overlaid with MEM (Thermo Fisher) with 8% FBS and 1.6% agarose. After 48 h, plates were stained with 0.05% neutral red (Sigma-Aldrich) and plaques were counted to calculate virus titers expressed as PFU/ ml.

### Histopathology

Lung tissues were inflated by injection of 1 ml 4% paraformaldehyde (PFA) into the left lung. Lungs were then fixed in 4% PFA for 3 days at 4°C and were submerged in 10% buffered formalin (Fisher Scientific, Waltham, MA, USA). Samples were sent to the Histopathology Laboratory Core at UTMB, where the paraffin-embedded tissue blocks and 10-μm tissue sections were prepared and hematoxylin and eosin (H & E) staining was performed.

## Statistical analysis

Data analysis for survival rate, viral load, cytokine production, antibody titers, and T cell responses was performed using GraphPad Prism software. Data were presented as means ± SEM. *P* values of these experiments were calculated with a non-paired Student’s t-test.

## Author contributions

A.A., C.L., M.J., B.H., and J.Z., performed the experiments. A.A., and T.W. designed the experiments. X.X., and P.W. provided critical reagents and supplies, A.A., C.L., M.J., B.H., J.Z., B.P., and T.W. analyzed the data. A.A., and T.W. wrote the initial draft of the manuscript and other coauthors provided editorial comments.

## Acknowledgements

This work was supported in part by NIH grants R01AI176670 (T.W.) and R01 NS125778 (T.W.). C.L. was a recipient of a summer internship from NIAID T35 training grant (T35AI078878, PI: T.W.).

## Competing interests

Authors declare that they have no competing interests.

## Supplementary Figure Legend

**Supplementary Figure 1. VSA-2-RBD and QS-21-RBD vaccination induced strong SARS-CoV-2 IgG binding antibodies in mice**.BALB/c mice were prime-boost immunized with mock (PBS), 10 or 15 µg of QS-21-RBD, or VSA-2-RBD via s.c. route. At 31 DPV, sera were collected. ELISA O.D. values of IgG against SARS-CoV-2 RBD were measured. n = 2 to 3.

## Notes

### Competing Interest Statement

The authors have declared no competing interest.

