## Supplementary figures and images for "VSA-2, a novel plant-derived adjuvant for SARS-CoV-2 subunit vaccine"

### Supplementary Fig 1

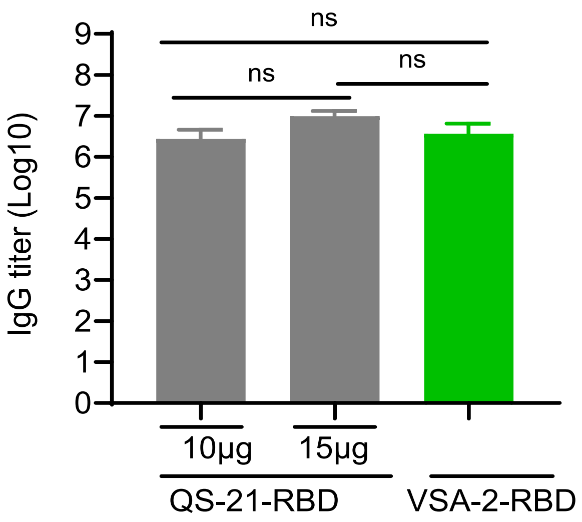


**Supplementary Figure 1**
